# The natural compound TMYX reduces SAN cells rate by antagonizing the cAMP modulation of f-channels

**DOI:** 10.1101/2021.11.16.468855

**Authors:** Chiara Piantoni, Manuel Paina, David Molla, Sheng Liu, Giorgia Bertoli, Hongmei Jiang, Yanyan Wang, Yi Wang, Yi Wang, Dario DiFrancesco, Andrea Barbuti, Annalisa Bucchi, Mirko Baruscotti

**Author notes:** Co-corresponding authors: Mirko Baruscotti, Università degli Studi di Milano, Department of Biosciences, via Celoria 26, 20133 Milano, Italy., Annalisa Bucchi, Università degli Studi di Milano, Department of Biosciences, via Celoria 26, 20133 Milano, Italy. These authors contributed equally to the paper. Institute of Neurophysiology, Hannover Medical School, Carl-Neuberg-Str.1, 30625 Hannover, Germany. AXXAM S.p.A., via Meucci 3, 20091 Bresso (Milano), Italy. **Author’s Contribution** C.P., M.P., A.Bu. and M.B.: performed statistical analysis; D.D.F, A.Bu., and M.B.: handled funding and supervision; C.P., M.P. and D.M.: acquired the data; C.P., M.P., D.D.F., S.L., W.Y., A.Bu. and M.B.: conceived and designed the research; C.P., M.P., A.Bu. and M.B.: drafted the manuscript; C.P., M.P., D.M., S.L., G.B., Y.W., W.Y., W.Y., H.J., A.Ba., D.D.F., A.Bu. and M.B.: made critical revision of the manuscript for key intellectual content. C.P. and M.P. contributed equally to this paper.

## Abstract

Tongmai Yangxin (TMYX), is a complex compound of a Traditional Chinese Medicine (TCM) used to treat several cardiac rhythm disorders; however, no information regarding its mechanism of action is available. In this study we provide a detailed characterization of the effects of TMYX on the electrical activity of pacemaker cells and unravel its mechanism of action.

Single-cell electrophysiology revealed that TMYX elicits a reversible and dose-dependent (2/6 mg/ml) slowing of spontaneous action potentials rate (−20.8/-50.2%) by a selective reduction of the diastolic phase (−50.1/-76.0%). This action is mediated by a negative shift of the I_f_ activation curve (−6.7/-11.9 mV) and is caused by a reduction of the cAMP-induced stimulation of pacemaker channels. We provide evidence that TMYX acts by directly antagonizes the cAMP-induced allosteric modulation of the pacemaker channels. Noticeably, this mechanism functionally resembles the pharmacological actions of muscarinic stimulation or β-blockers, but it does not require generalized changes in cytoplasmic cAMP levels thus ensuring a selective action on rate.

In agreement with a competitive inhibition mechanism, TMYX exerts its maximal antagonistic action at submaximal cAMP concentrations and then progressively becomes less effective thus ensuring a full contribution of I_f_ to pacemaker rate during high metabolic demand and sympathetic stimulation.

**Funding sources:** This work was supported by grants from Tianjin Zhongxin Pharmaceutical Group Co., Ltd. Le Ren Tang Pharmaceutical Factory P.R. China. The financial supporter played no role in the study design, data collection and analysis, decision to publish, or the preparation of the manuscript.

## Introduction

Traditional Chinese Medicine (TCM), one of the oldest organized healing systems in human history, is based on a holistic view of both the disease state and the associated therapy. For this reason, the perfect synthesis of TCM pharmacology is based on complex drugs composed by a mixture of different elements/herbs whose aim is to target the causes of the disease, modulate other aspects of the body wellness, and contrast toxicity (Chen et al., 2006). Western medicine follows a different perspective since it focuses on the molecular mechanisms and defines this level as its therapeutic target. Despite these differences, both approaches have reached reliable standards (Tang & Huang, 2013; Tu, 2016). These different views are now converging thanks to modern pharmacological and molecular studies whose approach is to experimentally challenge the efficacy of TCM drugs and to isolate active molecules that could represent novel acquisitions to the western pharmacopeia (Burashnikov, Petroski, Hu, Barajas-Martinez, & Antzelevitch, 2012; Chen et al., 2006; Efferth, Li, Konkimalla, & Kaina, 2007; Lao, Jiang, & Yan, 2009; Tang & Huang, 2013).

Based on these premises, we focused our study on Tongmai Yangxin (TMYX), a TCM botanical drug composed by at least 80 single molecular components among which flavonoids, coumarins, iridoid glycosides, saponins and lignans (Tao et al., 2015). In China, TMYX is used to treat several diseases including cardiovascular conditions (such as coronary artery disease, palpitation, heart failure, and angina). Interestingly, metabolomics analysis, carried out in a registered clinical trial on stable angina patients, has highlighted a reduction of serum markers of cardiac metabolic disorders, oxidative stress, and inflammation (Cai et al., 2018; Fan et al., 2016; Guo et al., 2020). In addition to this cardio-protective role, TMYX is also used as an antiarrhythmic agent (Cai et al., 2018; National Pharmacopeia Committee), but the mechanisms underlying this action are unknown. We therefore carried out *in-vitro* investigations in rabbit sinoatrial node (SAN) cells to verify the hypothesis that TMYX is also able to modulate their spontaneous activity. Our data confirm that TMYX reduces the rate of SAN cells by selectively reducing the slope of the diastolic depolarization. This action is similar to that of ivabradine which is a selective blocker of the pacemaker current (I_f_) and, at present, the only pure heart rate-lowering agent approved for clinical use in several western countries (Koruth, Lala, Pinney, Reddy, & Dukkipati, 2017). We have further discovered that TMYX modulates the whole-cell I_f_ current by inducing a cholinergic-like shift of the voltage-dependence of channels activation, and the underlying mechanism is a competitive antagonism of the cAMP-induced channel activation. Since the I_f_ current is a major contributor of the early part of the diastolic depolarization phase of pacemaker cells, its selective block is associated with bradycardia without negative inotropic effects (typical, for example, of β-blocker agents). Despite TMYX and ivabradine inhibit the I_f_ current with different molecular mechanisms, both these mechanisms functionally converge to a selective modulation of the early part of the diastolic depolarization of SAN cells and this raises a potential pharmacological interest in the active principle of this TCM drug.

## Results

We first investigated whether TMYX could modify the spontaneous electrical activity of rabbit SAN myocytes. In Figure 1A representative time-courses of Action Potential (AP) rate (top) and sample AP traces (bottom), recorded in the absence (Control) and in the presence of two different concentrations (2 and 6 mg/ml) of TMYX, are shown. TMYX caused a reversible and dose-dependent rate slowing (2 mg/ml: −20.8±1.6%, n=12 and 6 mg/ml: −50.2±6.5%, n=8 from mean control values of 3.6±0.1 Hz and 3.9±0.3 Hz, respectively). The analysis was then extended over a wider range of concentrations and the Hill fitting of the experimental dose-response data points distribution yielded a half-inhibitory value (k) of 4.9 mg/ml, a maximal block value (y_max_) of 92.7%, and a Hill coefficient (h) of 1.3 (Figure 1B). At the highest concentration tested (60 mg/ml), the average rate reduction was 86.7±6.3% (n=6); in 3 of these cells the activity was completely abolished. In all experiments, the effect of TMYX on rate was fully reversible after washout.

**Figure 1:**
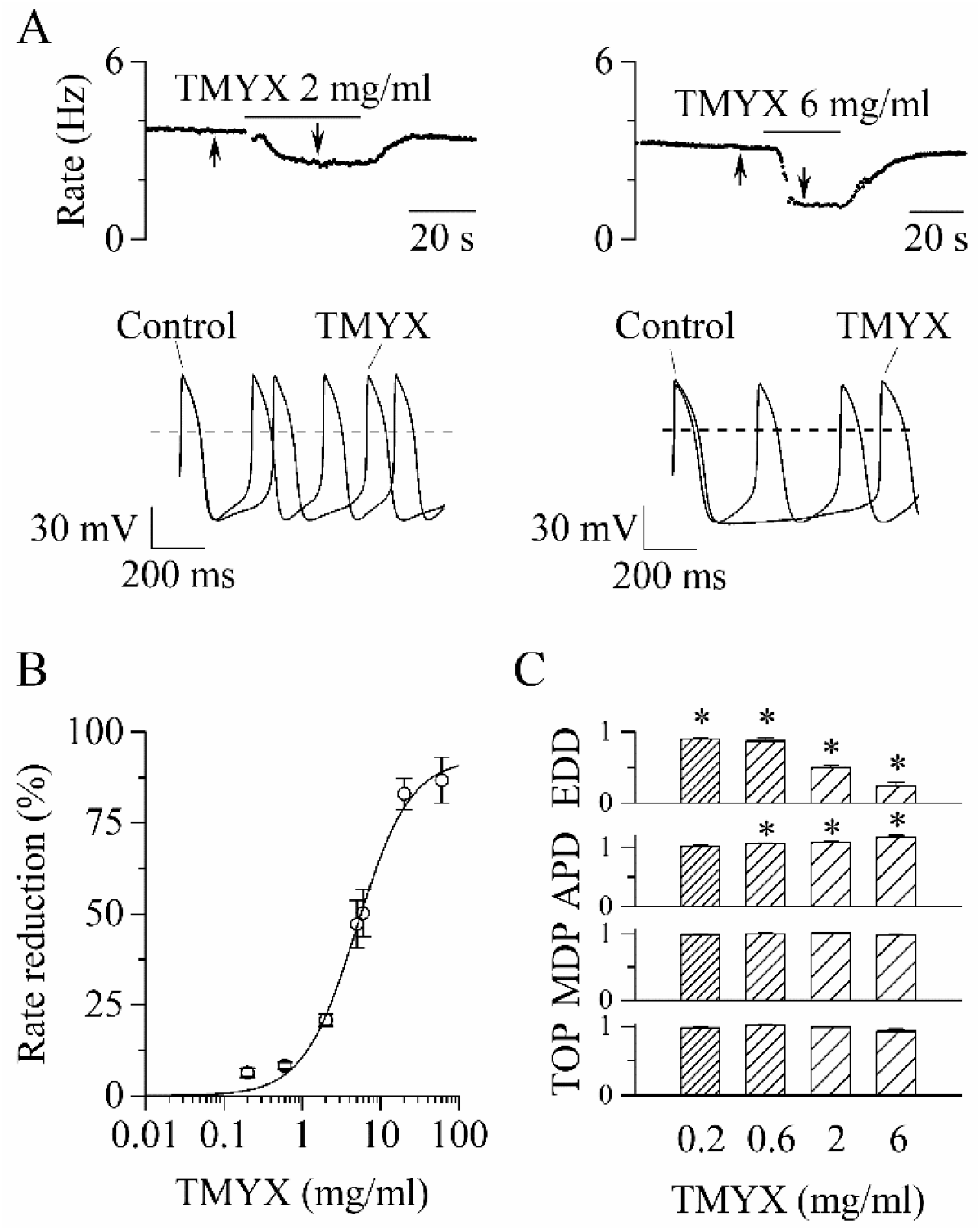
TMYX reduces the spontaneous rate of rabbit SAN myocytes. A: Representative time-courses (top) and sample traces (bottom) of spontaneous APs recorded from rabbit SAN cells in control conditions and in the presence of TMYX (2 and 6 mg/ml). Here and in other figures the arrows indicate the time of recording of the sample traces. B: Dose-response relationship of the AP rate reduction induced by TMYX; each point represents the mean±SEM% value obtained at the following doses: 0.2, 0.6, 2, 5, 6, 20, 60 mg/ml (n=68). The Hill fitting (full line, y=y_max_/(1+(k/x)^h^) yielded the following values: y_max_=92.7%, k=4.9 mg/ml, and h=1.3. C: Summary of the effects of TMYX on the AP parameters (n=7-12, details in the methods) normalized to the corresponding control values. Statistical analysis was carried out prior to normalization, *P<0.01 vs control (Student’s Pair t-Test).

To dissect the action of the drug during the various phases of the AP, we quantitatively evaluated specific AP parameters (Early Diastolic Depolarization, EDD; AP Duration, APD; Maximum Diastolic Potential, MDP; Take Off Potential, TOP) in the absence and during perfusion of different doses of TMYX (0.2, 0.6, 2, and 6 mg/ml). As shown in Figure 1B, the spontaneous rate was significantly reduced at all doses investigated, and this effect was for the largest part caused by a significant decrease of the EDD (rate: − 6.3±1.2%, −8.2±0.9%, −20.8±1.6%, −50.2±6.5%; EDD:-9.7±1.5%, −12.8±4.4%, −50.1±3.7%, − 76.0±5.7%). A small increase of the APD was observed at doses ≥0.6 mg/ml (6.2±0.8%, 8.7±1.1%, 17.1±3.7%); TOP and MDP were not affected.

Figure 1 provides evidence that TMYX lowers AP rate mainly by affecting the pacemaker mechanisms governing the EDD process. Since the I_f_ current is relevant to the generation of this phase (Bucchi, Baruscotti, Robinson, & DiFrancesco, 2007; DiFrancesco, 1993), we wondered whether this current could be a target of TMYX. We initially explored the effects of TMYX both on the voltage-dependence and on the maximal conductance of I_f_. To this aim we used a double-pulse protocol which allows to observe the effect of a drug on the current both near the half-activation voltage (−65 mV), and at full activation (−125 mV, Figure 2A). Perfusion of SAN cells with TMYX 6 mg/ml modified the current at both voltages, but in opposite directions: in the sample recordings shown in Figure 2A, at −65 mV the current was reduced by 40.5%, while at −125 mV was increased by 12.9%. This apparently paradoxical behavior was observed in all cells investigated (n=6 cells) and a possible explanation requires the combination of two contrasting effects: a negative shift of the activation curve and an increase of the maximal conductance. To evaluate this possibility, we carried out a quantitative characterization of these effects. The activation curves of the I_f_ current were measured in n=7 cells before and during TMYX (6 mg/ml) and mean±SEM values are plotted in Figure 2B. Boltzmann fitting of experimental data confirmed a significant hyperpolarizing shift of the activation curve (11.9 mV, P<0.01). If considered alone this effect would tend to decrease the contribution of the current to pacemaker depolarization, hence to rate slowing.

**Figure 2:**
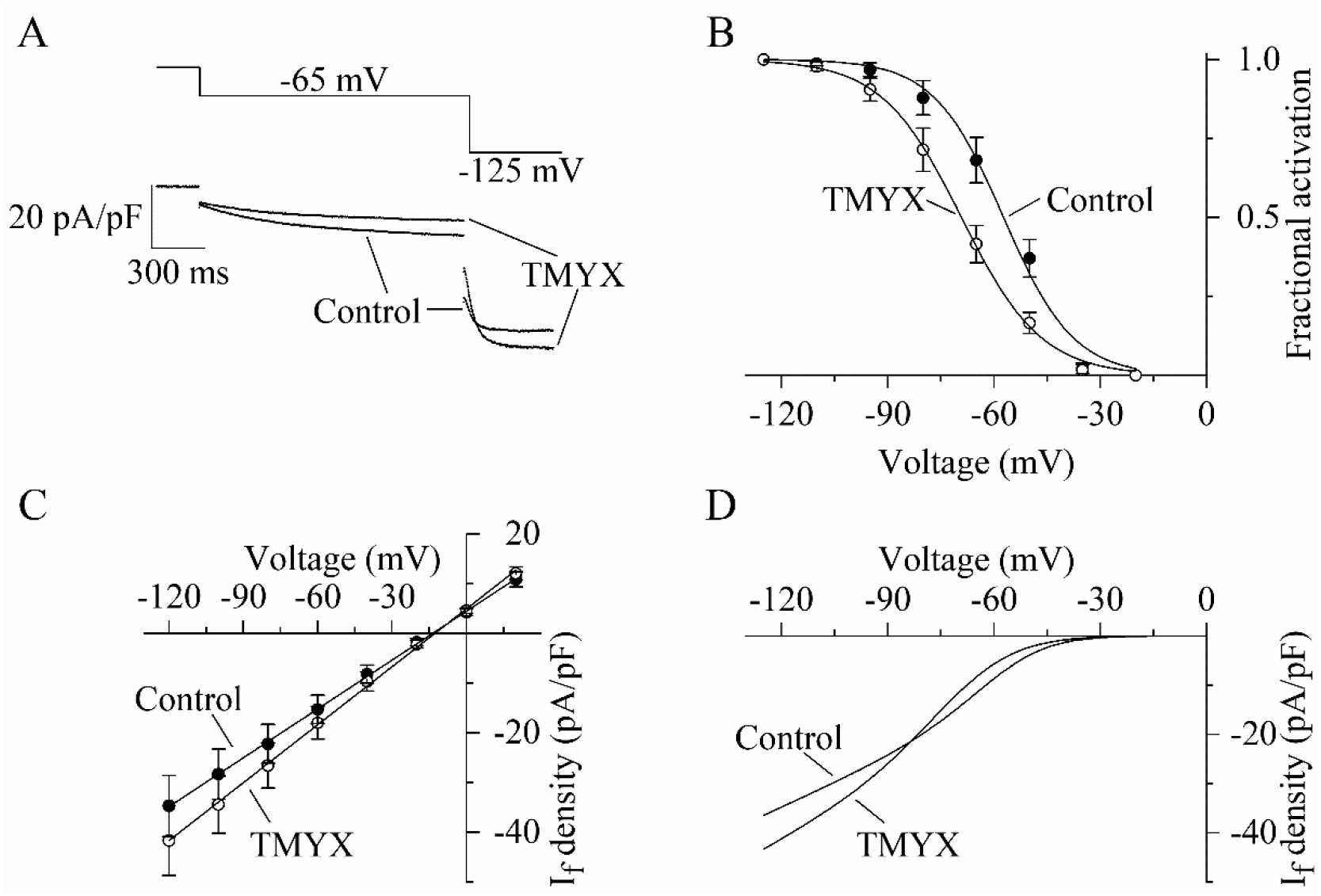
Dual action of TMYX on the voltage-dependence and maximal conductance of the I_f_ current. A: Representative whole-cell currents elicited by a double-step protocol (−65 mV/1.5 s and −125 mV/0.5 s; holding potential −35 mV) before (Control) and during drug (TMYX 6 mg/ml) perfusion. B: Voltage-dependent activation curves obtained in control conditions (filled circles) and during TMYX perfusion (empty circles). Boltzmann fitting (full lines, y=1/(1+exp((V-V_½_)/s)) of mean fractional activation values (n=7 cells) yielded the following half-activation (V_½_) and inverse-slope factors (s) parameters: − 57.3 mV and 9.8 mV (Control) and −69.2 mV and 11.3 mV (TMYX); the shift caused by TMYX is statistically significant (P<0.01, Extra sum-of-squares F test). C: Mean fully-activated current/voltage (I/V) relations measured before (filled circles) and during drug perfusion (empty circles, n=5 cells). Linear fitting yielded reversal potentials of −13.6 mV and −12.7 mV and slopes of 0.328 and 0.389 (pA/pF)/mV in control and in the presence of TMYX, respectively; the slopes are significantly different (P<0.01, linear regression analysis test). D: Steady-state I/V fitting curves obtained by multiplying the activation curves (Boltzmann fitting, panel B) and fully-activated I/V relation (linear fitting, panel C) in control condition and in the presence of the drug.

To better investigate the TMYX-induced current increase at −125 mV, we measured the fully-activated current/voltage (I/V) relation in control condition and in the presence of TMYX (6 mg/ml, n=5; Figure 2C). Linear fitting of mean±SEM data confirmed that TMYX increased the slope of the fully-activated I/V relation by 18.6%.

Data in Figure 2A,B,C thus indicate that TMYX exerts functionally opposite effects on the voltage-dependent availability of the current, which is decreased, and on the maximal conductance, which is increased. This observation is better illustrated by considering the steady-state I_f_ current curves (Figure 2D): at voltages more positive than the cross-over point (−83 mV) the prevalent effect of TMYX is a current reduction due to the leftward shift of its activation curve, while at more negative voltages the increase in conductance prevails. A similar effect was observed in the presence of a lower dose (2 mg/ml) of TMYX (Figure S1). To further strengthen this finding, a train of different activating steps was delivered prior to and during TMYX (6 mg/ml) exposure, and the mean±SEM steady-state current amplitudes (n=7 cells) are plotted in Figure S2. Fitting of experimental data with the following equation I_density_=(a*V+b)*(1/(1+exp((V-V_½_)/s))), which combines the linear I/V behavior with the Boltzmann sigmoidal voltage-dependence, confirmed the presence of a cross-over phenomenon in the steady-state I/V relations.

Taken together, data presented in Figures 2, Figures S1,S2 reveal that at diastolic voltages TMYX reduces the I_f_ contribution by shifting its voltage-dependence to more negative values. Since the cholinergic control of the I_f_ current, and thus of SAN rate, operates via a similar mechanism, we asked whether one or more components of TMYX could act as muscarinic agonist. To address this point, we compared the effects of TMYX (6 mg/ml) and Acetylcholine (ACh, 1 µM) on I_f_ and on cell rate, as measured in the absence and presence of the muscarinic blocker atropine (10 µM).

I_f_ current traces recorded at −65 mV in control and in the presence of either TMYX (top) or ACh (bottom) delivered alone (left) or in combination with atropine (right), are shown in Figure 3A.

**Figure 3:**
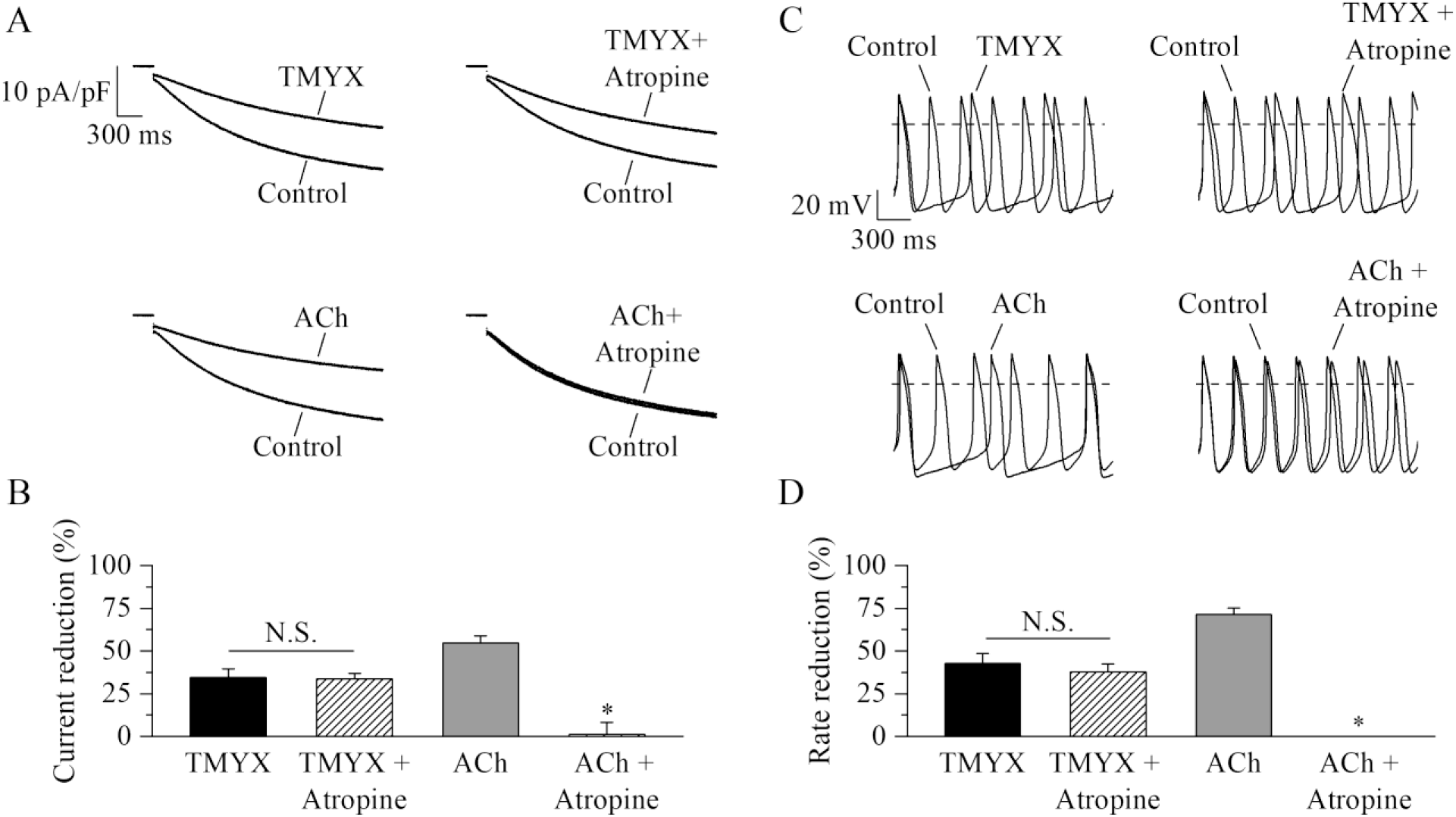
TMYX action does not involve muscarinic receptor activation. A: Representative sample current traces recorded during steps to −65 mV in the presence and in the absence of TMYX (6 mg/ml, top) and ACh (1 µM, bottom) delivered alone (left) or in combination with atropine (10 µM, right). B: Mean±SEM steady-state current reduction. Atropine did not modify the action of TMYX (TMYX, −34.6±4.9%; TMYX+Atropine, −33.5±3.4%, n=6) but abolished the effect of ACh (ACh, −54.7±4.0%; ACh+Atropine, 1.2± 6.9%, n=6). N.S. Not Significant, P=0.594; *P<0.01 *vs* ACh (Student’s Pair t-Test). C: Representative APs recorded in the same condition as in panel A. D: Mean±SEM rate reduction. Atropine did not reduce the ability of TMYX to induce cell bradycardia (TMYX, −42.6±5.8%; TMYX+Atropine, −37.6±4.8%, n=6), but abolished the action of ACh (71.5±3.6) (n=7). N.S. Not Significant, P=0.807; *P<0.01 *vs* ACh (Student’s Pair t-Test).

The TMYX-induced reduction of the I_f_ current was not modified by atropine (TMYX:-34.6±4.9%, TMYX+Atropine:-33.5±3.4%, n=7), while atropine abolished the effect of ACh (n=6; Figure 3A,B). In line with the findings on I_f_, we also observed that the muscarinic block did not antagonize the rate-slowing effect elicited by TMYX on SAN cells (TMYX:-42.6±5.8%; TMYX+Atropine:-37.6±4.8%, n=6; Figure 3C,D). On the other hand atropine abolished the ACh-induced rate slowing (n=7; Figure 3C,D).

We also asked whether TMYX could exert its inhibitory action by interfering with adenosine, a well-known modulator of I_f_ whose action is based on a negative shift of the activation curve (Zaza, Rocchetti, & DiFrancesco, 1996). Data shown in Figure S3 also demonstrate that the adenosine receptor is not involved in the TMYX-induced inhibition of the current.

After excluding the involvement of a direct activation of the muscarinic (and adenosine) receptors, we next asked whether TMYX could exert its effect on the I_f_ activation curve by interfering with a downstream effector, and particularly on the cAMP-dependent modulation of the channel. This hypothesis was based on the well-established evidence that the voltage-dependent availability of I_f_ is controlled by the binding/unbinding of cAMP molecules to the pacemaker f-channels (DiFrancesco & Tortora, 1991; James & Zagotta, 2018).

To investigate the existence of a possible functional interference between cAMP and TMYX, we performed the experiments shown in Figure 4, where the effect of TMYX (6 mg/ml) on the whole-cell I_f_, measured in the diastolic range of potentials, was assessed in the absence (Basal) and in the presence of two concentrations of cAMP in the pipette solution (10, 100 µM).

**Figure 4:**
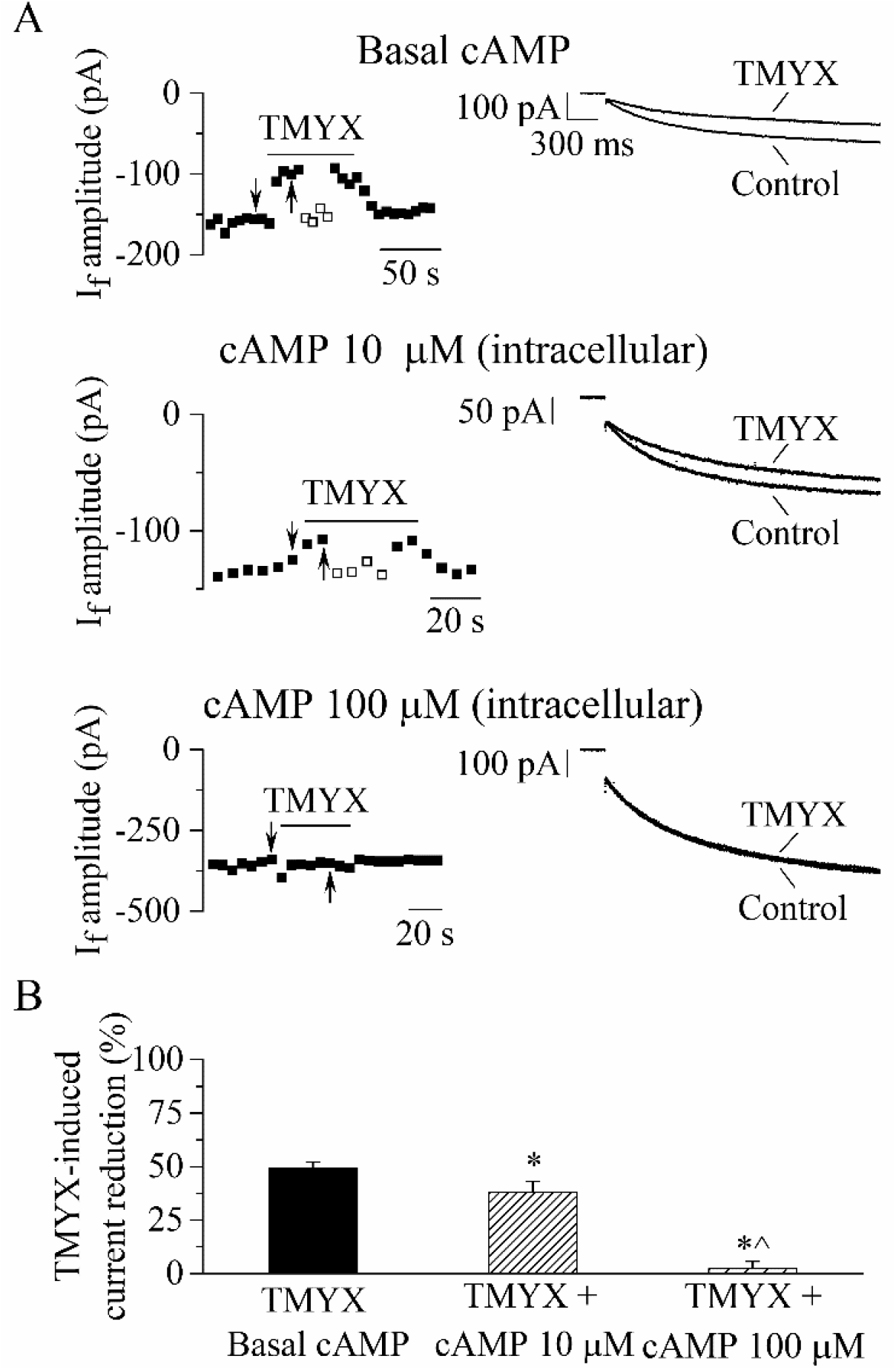
The TMYX-induced reduction of the whole-cell I_f_ is counteracted by increasing concentrations of intracellular cAMP. A: Representative time-courses (left) of the steady-state amplitudes of the current recorded at −65 mV/2.75 s (sample traces, right) in control conditions and in the presence of TMYX (6 mg/ml). Experiments were carried out in the absence (top, n=15), and in the presence of 10 µM (middle, n=6) and 100 µM (bottom, n=6) cAMP in the pipette intracellular solution. Empty squares in top and middle panels indicate steady-state currents recorded after manual adjustment of the holding level (ΔV) to compensate for the inhibitory effect induced by TMYX. B: Bar-graph of the TMYX-induced current reductions (mean±SEM%) obtained in the three different conditions: TMYX/Basal cAMP, −49.1±3.0%, n=15; TMYX+cAMP 10 µM, −37.9±5.1%, n=6; TMYX+cAMP 100 µM, −2.3±3.3%, n=6. *P=0.049 cAMP 10 µM *vs* Basal cAMP; *P<0.01 cAMP 100 µM *vs* Basal cAMP; ^P<0.01 cAMP 100 µM *vs* cAMP 10 µM (One-Way ANOVA followed by Fisher’s LSD post-hoc test multiple comparisons).

Representative time-courses (left) and current traces (right), recorded in the three different experimental conditions during repetitive hyperpolarizing steps to −65 mV in the absence and presence of TMYX, are presented in Figure 4A. A progressive loss of modulatory efficacy of TMYX clearly appears as the intracellular cAMP content increases. As shown in the bar-graph plots in Figure 4B, a quantitative evaluation of the results yielded the following TMYX-induced current reductions (mean±SEM): Basal cAMP, −49.1±3.0%, n=15; cAMP 10 µM, −37.9±5.1%, n=6; cAMP 100 µM, −2.3±3.3% (n=6; all conditions are significantly different, see legend). The modulatory efficacy of the drug, and its dependence upon intracellular cAMP, was also estimated by means of the ΔV method (empty squares in Figure 4A; Material and Methods for details) since this analysis allows to assess the shift of the I_f_ activation curve (Accili & DiFrancesco, 1996; DiFrancesco, Ducouret, & Robinson, 1989). Mean±SEM TMYX-induced hyperpolarizing ΔV (shift) values were: Basal cAMP, 6.3±0.3 mV, n=15; cAMP 10 µM, 4.5±0.7mV, n=6; cAMP 100 µM, 0.45±0.45 mV (n=6; all significantly different, P<0.01, One-Way ANOVA followed by Fisher’s LSD post-hoc test multiple comparisons). However, since, in addition to its effect on the activation curve, TMYX also affects the maximal conductance of the current, the ΔV values measured in the experimental paradigm of Figure 4 represent an underestimation of the absolute shift.

The evidence that the modulatory efficacy of TMYX is counteracted by increasing concentrations of cAMP suggests the intriguing hypothesis of an antagonistic action between these two compounds; additional evidence supporting a mutual interference is presented in Figure S4. In this case the cAMP content of SAN cells was experimentally raised by: *i*) inhibiting its degradation using a phosphodiesterase inhibitor (IBMX, 100 µM) and *ii*) favoring its overproduction using an activator of the Adenylyl Cyclase (Forskolin, 100 µM). The ability of TMYX (6 mg/ml) to reduce the I_f_ current was then quantified in the presence of different combinations of these substances and of cAMP (10 µM). The bar-graphs shown in Figure S4A,B confirm the presence of an inverse dependence between cAMP levels and TMYX efficacy. However, these experiments do not provide details on the underlying mechanism.

We therefore proceeded by taking advantage of the inside-out macro-patch configuration since this experimental approach allows testing whether TMYX has a membrane-delimited effect (by directly acting on f-channels) or requires instead the involvement of cytoplasmic elements controlling cAMP production and degradation. In Figure S5A, the I_f_ current was elicited by a train of hyperpolarizing steps to −105 mV to test the effect of TMYX (6 mg/ml) delivered in the absence of cAMP; no modulation of the current was ever observed (n=4 patches, P=0.125, Student’s Pair t-Test). An analysis extended to a wider range of voltages is presented in Figure S5B,C where a slowly-activating voltage ramp (−35/-145mV) was employed to measure the steady-state I/V curves (panel B) and the associated conductance/voltage (g/V) curve (panel C), in the absence (Control) and in the presence of TMYX (6 mg/ml). Statistical analysis revealed that neither half-activation (V_½_) nor maximal conductance (g_max_) parameters were affected by the presence of TMYX (Figure S5D). This evidence demonstrates that TMYX does not influence the intrinsic properties of f-channels and raises the possibility that its inhibitory effect can only occur in the presence of a concurrent cAMP-dependent modulation of the channels.

To verify this possibility we first evaluated the shift of the I_f_ voltage dependence induced by cAMP using the ΔV method (Accili & DiFrancesco, 1996; DiFrancesco et al., 1989) and then the ability of TMYX to reverse this shift (Figure 5). Inside-out I_f_ currents were elicited by a train of hyperpolarizing steps (−105 mV) while membrane patches were exposed to different cAMP concentrations (1, 10, 100 µM) delivered alone and in the presence of TMYX (6 mg/ml). Representative time-courses, current traces, and the corresponding analysis are shown in Figure 5. Exposure to cAMP elicited a dose-dependent increase of the current, which was quantified as the voltage-correction necessary to restore steady-state control current levels (ΔV_cAMP/Cont_: 6.1±0.5 mV, 12.8±0.5 mV, and 13.6±0.9 mV for 1, 10, 100 µM cAMP, respectively; Figure 5A, empty triangles). Addition of TMYX (cAMP+TMYX) resulted in a reversible reduction of cAMP action quantified as the ΔV correction required to compensate for the effect of TMYX (ΔV_TMYX/cAMP_: 3.5±0.4 mV, 4.7±0.6 mV, and 0 mV for 1, 10, 100 µM cAMP, respectively; Figure 5A, empty squares). The difference between experimental ΔV_cAMP/Cont_ and ΔV_TMYX/cAMP_ values represents the cAMP-induced shift in the presence of TMYX (ΔV_(cAMP+TMYX)/Cont_). Dose-dependent ΔV_cAMP/Cont_ (empty triangles) and ΔV_(cAMP+TMYX)/Cont_ (empty circles) values calculated for each patch are plotted in the left panel of Figure 5B, and Hill fittings of data points yielded half-maximal concentrations (k) of 1.17 µM and 5.66 µM for the two conditions, respectively. To better illustrate the antagonism exerted by TMYX (6 mg/ml) on cAMP, we calculated the TMYX-induced fractional inhibition by normalizing the TMYX-induced inhibition of cAMP action (ΔV_TMYX/cAMP_) to the corresponding full cAMP modulation (ΔV_cAMP/Cont_). This procedure was applied both on experimental data points and on the corresponding Hill fittings shown in Figure 5B,left and results are plotted in Figure 5B,right (filled diamonds and dashed line). This distribution demonstrates that in inside-out conditions TMYX antagonizes the effect of cAMP at intermediate (1, 10 µM), but not at high cAMP doses, and the maximal antagonistic effect (63.7%) was observed at a cAMP concentration of 0.8 µM.

**Figure 5:**
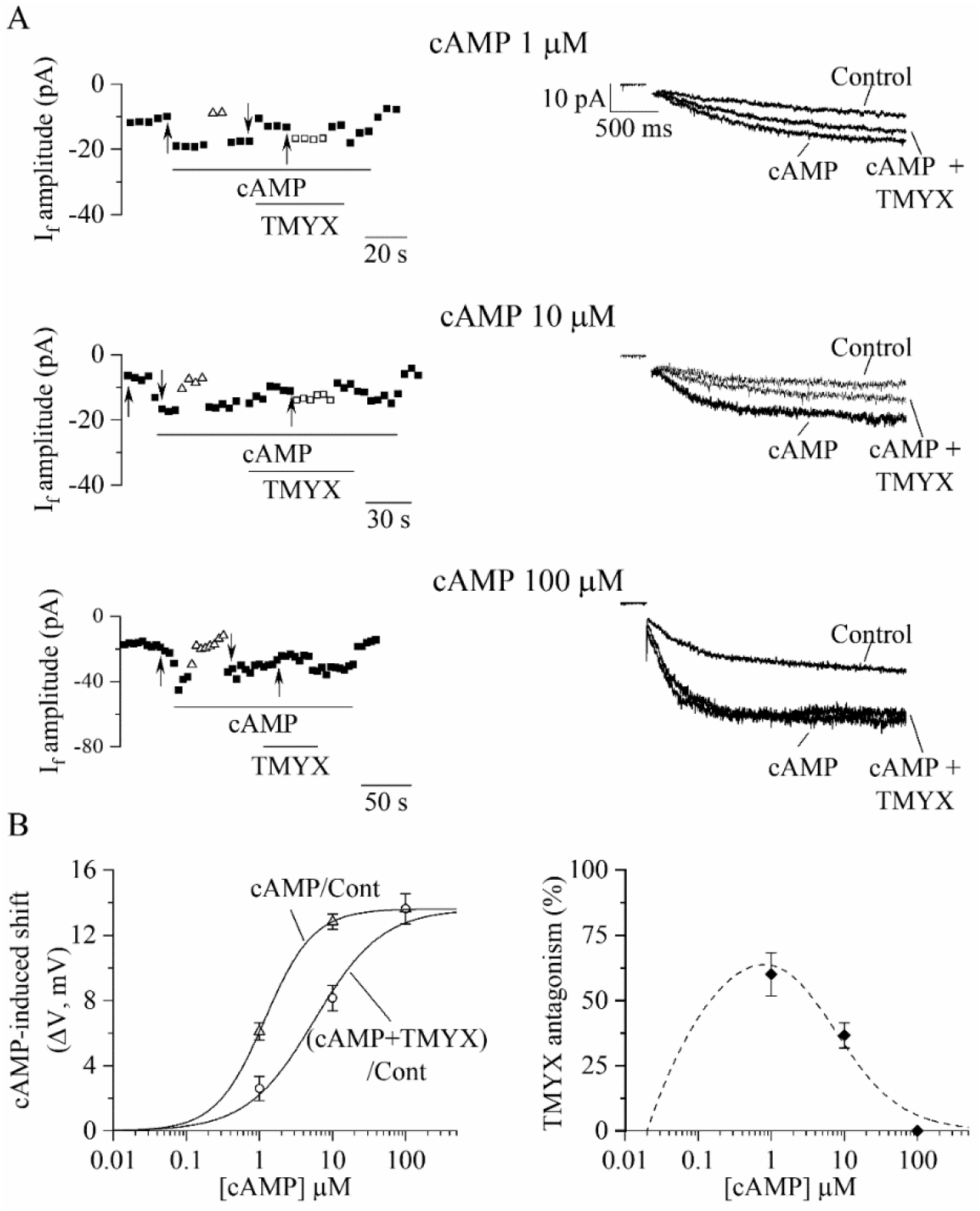
TMYX reduces the I_f_ current by antagonizing its cAMP-induced modulation. A: Sample time-courses (left) and current traces (right) of I_f_ amplitudes recorded in inside-out macro-patches during hyperpolarizing steps (−105 mV); cAMP (1, 10, 100 µM, n=4-6) was perfused alone (cAMP) or in combination with a fixed dose of TMYX (6 mg/ml, cAMP+TMYX). Empty triangles and squares represent current amplitudes observed after correcting the applied voltage (ΔV) to compensate for (and evaluate) both the effect of cAMP (triangles) and the ability of TMYX to reduce cAMP modulation (squares). B, left: cAMP-induced shifts of the I_f_ activation curve obtained in the presence of cAMP alone (cAMP/Cont, empty triangles) and of cAMP+TMYX ((cAMP+TMYX)/Cont, empty circles). The continuous lines represent dose-response Hill fittings of experimental data-points (cAMP/Cont: k=1.17 µM and h=1.30; (cAMP+TMYX)/Cont: k=5.66 µM and h=0.94; 13.6 mV was the maximal shift experimentally measured and was therefore taken as y_max_ for both conditions). B, right: the antagonism exerted by TMYX on cAMP action was calculated as the fractional inhibition of the I_f_ current as derived from the Hill fittings (Hill_cAMP/Cont_-Hill_(cAMP+TMYX)/Cont_)/Hill_cAMP/Cont_ and from experimental points (diamond symbols, see text for details).

## Discussion

Natural botanical compounds commonly used in TCM have recently become of interest also to modern pharmacological studies whose approach is to scientifically challenge their efficacy and to isolate active molecules that could represent novel acquisitions to the western pharmacopeia (Chen et al., 2006; Tang & Huang, 2013; Tu, 2016). Indeed, several studies have demonstrated the safe and beneficial effects of TCM drugs on different pathologies including cancer and cardiovascular diseases (Efferth et al., 2007; Li et al., 2013; Pommier, 2006). Interestingly, cardiovascular TCM drugs often target ion channels; for example, the antiarrhythmic agent Wenxin Keli binds to atrial Na^+^ channels according to a mechanism of potential relevance in the treatment of atrial fibrillation (D. Hu et al., 2016).

In this study we have characterized the effects of TMYX on the properties of pacemaker cells, since this drug is used in TCM to treat cardiovascular diseases including cardiac arrhythmias, Coronary Artery Disease (CAD) and angina (Cai et al., 2018; Fan et al., 2016).

### At low doses TMYX mainly controls the EDD and the I_f_ current

Our study indicates a reversible and dose-dependent depression of SAN cell rate due to a robust reduction of the slope of the EDD and by a limited prolongation of the APD (Figure 1). These actions are similar to those elicited by the selective I_f_ blocker ivabradine, which is the only pure heart rate-reducing drug used in western medicine for the treatment of angina and heart failure (Borer et al., 2012; Bucchi, Baruscotti, & DiFrancesco, 2002; Tardif, Ponikowski, Kahan, & Investigators, 2013; Thollon et al., 1997). When tested in SAN cells, TMYX (2 mg/ml, Figure 1B) and ivabradine (3 µM (Bucchi et al., 2007)) slow cell rate by 20.8% and 16.2% and prolong the APD by 8.6% and 9.4%, respectively. Similar effects of ivabradine (3 µM) have also been reported in SAN tissue preparation (rate: −19.6/-23.8%, APD50: +6.7/+8.9% (Thollon et al., 1997; Thollon et al., 1994)). Since the APD prolongation represents a pro-arrhythmic effect, the evidence that, at least in single SAN cells, ivabradine and TMYX act similarly on this parameter, suggests a dose-dependent safety of the drug in relation to AP prolongation-dependent arrhythmias. This observation correlates with the use of this drug in TCM clinic.

TMYX exerts a dual action on I_f_: a negative shift of the voltage-dependence and an increase of the maximal conductance, and the former action prevails at physiological voltages (Figure 2, Figures S1,S2). Interestingly, a negative shift of the activation curve is also the main mechanism during a moderate muscarinic stimulation (DiFrancesco et al., 1989); however, this mechanism is not shared with ivabradine. In SAN cells, a moderate cholinergic activation causes a reduction of cell cAMP content and this associates with a decreased cAMP-dependent modulation of sinoatrial HCN/funny channels; the opposite sequence of events occurs during adrenergic modulation of pacemaker rate (DiFrancesco, 1993). For this reason, the shift of the I_f_ voltage-dependence can be considered a readout parameter of the functional interaction between cAMP and HCN/funny channels.

### TMYX exerts a direct competitive antagonism on the cAMP-induced activation of the I_f_ current

Whole-cell experiments presented in Figure 2B reveal that, the shift induced by TMYX (6 mg/ml) is −11.9 mV, a value similar to the maximal shift induced by ACh (1 µM, 12.6 mV, (Accili, Robinson, & DiFrancesco, 1997)). This comparison thus suggests that the voltage dependent modulation of I_f_ induced by 6 mg/ml TMYX should approximate saturation. However, despite a similar effect on the current, 1 µM ACh blocks the spontaneous activity of SAN cells (DiFrancesco et al., 1989), while 6 mg/ml TMYX reduces rate only by ∼50% (Figure 1). Since TMYX does not act on the muscarinic receptor (Figure 3), this difference likely arises from the robust cholinergic activation of IK(_ACh)_. While TMYX does not activate the muscarinic receptor, it is conceivable that it may interfere with the cAMP-dependent modulation of the I_f_ current somewhere along the pathway downstream the receptor. This conclusion is further supported by the evidence that TMYX efficacy is independent from the stimulation of the adenosine receptor (Figure S3), whose effect on the I_f_ current is also mediated by a reduction of the cellular cAMP (Zaza et al., 1996).

The observation that the inhibitory action of TMYX on the whole-cell I_f_ is counteracted by increasing concentrations of intracellular cAMP (Figure 4, Figure S4), suggested a functional competitive antagonism. This hypothesis was further corroborated by the inside-out experiments (Figure 5, Figure S5) which revealed that TMYX does not act in the absence of cAMP. Several studies have shown that cAMP binding to the C-terminus of HCN channels initiates a domino-like structural rearrangements leading to the removal of the auto-inhibitory condition which is a hallmark of the cAMP-unbound HCN channels (Wainger, DeGennaro, Santoro, Siegelbaum, & Tibbs, 2001). The functional aspect of these events is an allosteric-driven shift of the open-close equilibrium towards the open state (Akimoto, VanSchouwen, & Melacini, 2018; DiFrancesco, 1999; Wainger et al., 2001). Our data indicate that TMYX exerts its competitive antagonism either by reducing the channel affinity for cAMP or by interrupting the structural relaxation. The competitive antagonism is clearly illustrated in Figure 5B,left where the comparison of the ΔV_cAMP/Cont_ and ΔV_(cAMP+TMYX)/Cont_ dose-response curves display the hallmarks of allosteric inhibition according to the concerted-symmetry model (I.H., 1975): a similar saturating effect (y_max_), a decrease in the half-maximal shifts (k), and a lower Hill coefficient (h). Furthermore, a dissociation constant (k_i_) value of 1.56 mg/ml was obtained for TMYX by applying the Schild equation (k_(cAMP+TMYX)/Cont_/k_cAMP/Cont_=1+[TMYX]/k_i_: where k_(cAMP+TMYX)/Cont_ and k_cAMP/Cont_ are half-maximal cAMP-induced shifts in the presence/absence of 6 mg/ml TMYX). This dose is compatible both with the half-inhibitory value observed for the TMYX action on rate (Figure 1B) and with the nearly maximal effect on I_f_ reported for the dose of 6 mg/ml (Figure 2B and previous comments).

Interestingly, the neuronal accessory protein TRIP8b modulates cAMP-dependence of HCN channels with a mechanism similar to that of TMYX. For this reason TRIP8b has raised interest since it may represent a therapeutic target for major depressive disorders (L. Hu et al., 2013; Lyman, Han, & Chetkovich, 2017; Saponaro et al., 2014).

An important parallelism exists between the actions of β-blockers and TMYX since both reduce the cAMP-induced activation of f-channels: TMYX antagonizes the action of cAMP directly at the channel level (Figure 5 and Figure S5), while β-blockers inhibit the β-receptor-cascade and the associated cAMP synthesis. Despite these different mechanisms, the common functional outcome is the modulation of the I_f_ activation curve (Figures 2, 5 and Figure S2). According to the mechanism of action identified in our study, the putative active molecule of TMYX directly regulates the pacemaker f-channel but does not modulate the overall cAMP content of the cell and, for this reason, it is expected to have a selective action on chronotropic control of rate without affecting the inotropism. Multiple effects are instead associated with β-block since a reduction of cell cAMP necessarily affects other processes such as the PKA modulation of other ion channels. For this reason, the identification of novel pure bradycardic agents is an important pharmacological aim (Nikolovska Vukadinovic et al., 2017).

Although robust experimental data on the effects of therapeutic use of TMYX are not yet available, the mechanism of action is compatible with its use in the treatment of CAD and irregular heart beat (Fan et al., 2016). In the inside-out configuration, the maximal antagonistic effect (63.7%) of TMYX at the dose of 6 mg/ml is observed at a cAMP concentration of 0.8 µM and progressively decreases at higher doses (Figure 5B,right). This behavior reveals an additional well-suited physiological and pharmacological feature since it allows recruiting full I_f_ current and rate modulation when tachycardic stimuli (cAMP levels) are boosted to maximum. The rationale behind clinical use of TMYX in TCM is further supported by the evidence that a synthetic derivative of TRIP8b can prevent the β-adrenergic control of SAN cell rate and of I_f_ (Saponaro et al., 2018) and this effect represents proof of principle for further studies and development in applied pharmacology (Proenza, 2018).

Finally, it should be mentioned that TMYX is also used to treat Premature Ventricular Complexes (PVCs; personal communication to MB) and, in some cases, PVCs are associated with adrenergic stimuli, high cAMP cell content, and expression of f-channels (Cantillon, 2013; Lee, Walters, Gerstenfeld, & Haqqani, 2019; Oshita et al., 2015). The evidence that ivabradine may prevent this ectopic activity (Kuwabara et al., 2013) further supports a causative association, and allows to speculate that TMYX could in principle have a therapeutic role.

In conclusion, TMYX slows spontaneous rate of SAN cells and the underlying mechanism is a selective depression of the diastolic depolarization operated through an antagonistic action on cAMP-induced pacemaker channel activation. Comparison with other pharmacological chronotropic modulators (ivabradine and β-blockers) reveals that TMYX may have an interesting and safe profile. In addition, as pointed out by Akimoto et al. (Akimoto et al., 2018), targeting the cAMP binding domain may represent an interesting future perspective for selective modulation of HCN channels since it will reduce the possibility of unspecific interference with other channels. Although TMYX is composed by several components, its mechanism is compatible with the action of a single molecule; we therefore believe that future investigations should focus on this search in addition to provide exhaustive clinical data on TCM patients.

## Materials and Methods

All animal procedures performed in this study were carried out in accordance with the guidelines of the care and use of laboratory animals established by the Italian and UE laws (D. Lgs n° 2014/26, 2010/63/UE); the experimental protocols were approved by the Animal Welfare Committee of the Università degli Studi di Milano and by the Italian Ministry of Health (protocol number 1127-2015).

### Animal procedures and cell isolation

New Zealand female rabbits (0.8-1.2 kg) were anesthetized by intramuscular injection of xilazine (5 mg/kg) and euthanized by an overdose i.v. injection of sodium thiopental (60 mg/kg). The hearts were then quickly removed and placed in pre-warmed (37°C) normal Tyrode’s solution (mM: NaCl, 140; KCl, 5.4; CaCl_2_, 1.8; MgCl_2_, 1; D-glucose, 5.5; Hepes-NaOH, 5; pH 7.4) containing heparin (10 U/ml). After surgical isolation, the sinoatrial node was cut into 5-6 pieces and treated according to a standard procedure to obtain isolated SAN cells (Bucchi et al., 2007). Cells were kept alive and in optimal conditions at 4°C and used for electrophysiological recordings within 48 hrs.

### Experimental solutions

Spontaneous action potentials were recorded from single cells or small beating aggregates; during these recordings the cells were perfused with a normal Tyrode’s solution and the patch pipettes were filled with (mM): NaCl, 10; K-aspartate, 130; ATP (Na-salt), 2; MgCl_2_, 2; CaCl_2_, 2; EGTA-KOH, 5; Hepes-KOH, 10; creatine phosphate, 5; GTP (Na-salt), 0.1; pH 7.2. Similar solutions were used to record the I_f_ current in whole-cell condition with the addition of BaCl_2_ (1 mM) and MnCl_2_ (2 mM) to the extracellular Tyrode’s to block contaminating K^+^ and Ca^2+^ currents. In inside-out recordings the control solution used to perfuse the intracellular side of the excised patches contained (mM): NaCl, 10; K-aspartate, 130; CaCl_2_, 2; EGTA-KOH, 5; Hepes-KOH, 10; pH 7.2, and the patch-pipette solution contained (mM): NaCl, 70; KCl, 70; CaCl_2_, 1.8; MgCl_2_, 1; BaCl_2_, 1; MnCl_2_, 2; Hepes-NaOH, 5; pH 7.4. The resistance of patch pipettes used in whole-cell experiments measured 3–5 MΩ; larger pipettes (0.5–2 MΩ) were used during inside-out macropatch recordings.

Tongmai Yangxin was kindly provided by Le Ren Tang Pharmaceutical Factory (Tianjin, P.R. China) as a dry powder. A stock solution was daily prepared by dissolving the appropriate amount of substance in water (∼80 °C for 15 minutes); this solution was then filtered (pore size, 0.45 µm) to remove undissolved components. The stock solution was used to prepare the test solutions at the desired concentrations. Acetylcholine (ACh), atropine, cyclic adenosine monophosphate (cAMP), forskolin, 1,3-dipropyl-8-cyclopentylxanthine (DPCPX), and 3-isobutyl-1-methylxanthine (IBMX) were purchased from Sigma-Aldrich Corporation and used at the concentrations indicated in the text.

Control and test solutions were delivered to the cells through a fast perfusion system or loaded in the whole-cell pipette solution as indicated in the text.

### Patch-clamp experiments and data analysis

Experiments were carried out using the patch-clamp amplifier Axopatch 200B and the pClamp 10.7 software (Molecular Devices, CA); data were analyzed with Clampfit, OriginPro 2020 (Origin Lab, Northampton, MA), Prism 5 (GraphPad Softwares, CA), and a customized software.

Action Potentials (AP) were recorded from single cells or small beating aggregates and acquired at a sampling rate of 1-2 kHz. After acquisition, action potential traces were digitally smoothed by a 10-point adjacent averaging smoothing procedure and the time-derivative calculated according to a second polynomial, 8-point smoothing differentiating routine. AP traces were then processed with customized software to calculate the following parameters: Rate (Hz), Maximum Diastolic Potential (MDP, the most negative potential for each action potential), Take-Off Potential (TOP, the voltage at which the voltage derivative overtakes a fixed threshold of 0.5 mV/ms), Early Diastolic Depolarization (EDD, defined as the mean slope in the first half of the diastolic depolarization), Action Potential Duration (APD, the time interval between TOP and the following MDP); additional details can be found in (Bucchi et al., 2007). Experimental dose-response points presented in Figures 1B and 5B were fitted to the Hill equation (y=y_max_/(1+(k/x)^h^), where y_max_ is the maximal effect, x is the drug concentration, k is the drug concentration eliciting half-maximal block (Figure 1B) or half maximal shift (Figure 5B) and h is the Hill factor.

The activation curves of the whole-cell I_f_ current were obtained by applying a train of two consecutive voltage steps: the first pulse was delivered at test potentials (from −20 to −125 mV, increment between steps: −15 mV) to attain steady-state current activation, while the second step was delivered at −125 mV to ensure maximal activation. Normalized tail currents amplitudes at −125 mV represent the activation variable at each test potential. Mean±SEM fractional activation values, measured in control condition and in the presence of different concentrations of TMYX, were interpolated by the Boltzmann distribution (y=1/(1+exp((V-V_½_)/s)), where y is the fractional activation, V is voltage, V_½_ is the half-activation voltage, and s is the inverse-slope factor.

The fully-activated current/voltage (I/V) relations in whole-cell condition were obtained by applying a voltage protocol consisting of two sequential pulses: the cell was first hyperpolarized to −125 mV and then depolarized to different test potentials in the range −120/+20 mV (increment: 20 mV). After leakage correction the tail currents were normalized and plotted as a function of tail step voltages. Mean±SEM experimental values were interpolated by a linear fit (I_density_=(a*V+b)) to yield the fully-activated (I/V) curves.

The steady-state current/voltage (I/V) curves shown in Figure 2D, Figure S1D were obtained by multiplying the Boltzmann fitting of fractional activation and the linear fitting of the fully-activated I/V curve.

During macropatch inside-out experiments shown in Figure 5 the I_f_ current amplitude was elicited by hyperpolarizing steps to −105 mV from a holding potential of −35 mV. Steady-state I-V relations (Figure S5B) were recorded by means of hyperpolarizing ramps from −35 to −145 mV at a rate of −110 mV/min. Inside-out activation curves (Figure S5C) were derived from the steady-state I-V currents (see (Bois, Renaudon, Baruscotti, Lenfant, & DiFrancesco, 1997) for details) and fitted to Boltzmann equation.

In some analysis (Figures 4 and 5) the effect of TMYX on the I_f_ current was assessed by means of the ΔV method. This method quantifies the manual voltage adjustment (ΔV) of the holding potential which is introduced during drug delivery to compensate for the drug-induced current reduction and restore a steady-state current amplitude as the one observed prior to drug delivery (that is in control condition). The ΔV parameter measured in this condition was originally used by several authors (Accili & DiFrancesco, 1996; Altomare et al., 2003; Bois et al., 1997) to evaluate the shift of the activation curve induced by modulatory agents. In whole-cell experiments presented in Figure 4, however, this parameter (see text) is an underestimation of the shift of the activation curve since the ΔV adjustment must also compensate for the TMYX-induced increase of channel conductance. This limitation does not apply to data presented in Figure 5 since in the inside-out configuration (Figure S5B) we could not measure any increase of the channel conductance.

Whole-cell and inside-out experiments were carried out at 35±0.5 °C and at room temperature, respectively.

### Statistical analysis

No statistical method was used to predetermine sample size, but our samples sizes are similar to those reported in previous studies (Altomare, Tognati, Bescond, Ferroni, & Baruscotti, 2006; Bucchi et al., 2007; Milanesi, Baruscotti, Gnecchi-Ruscone, & DiFrancesco, 2006; Thollon et al., 1994; Van Bogaert & Pittoors, 2003).

All data are presented as mean±SEM values. Group comparisons were analyzed for statistical significance using Student’s Pair t-Test (correlated samples, Figures 1C, 3B,D and Figures S3B, S5A,D) or One-Way ANOVA followed by Fisher’s LSD post-hoc test (multiple comparisons, Figure 4B and Figure S4A,B).

Activation curves were compared using the Extra sum-of-squares F test, while the slopes of current/voltage (I/V) relations were evaluated through the linear regression analysis test (Figure 2B,C and Figure S1B,C).

Computational and statistical analysis were carried out with OriginPro 2020, OriginLab Corporation, USA and GraphPad Prism 5, GraphPad Software, USA.

Statistical significance is indicated by P values<0.05. Exact P values are provided except when P<0.01.

## Abbreviations

TCM: Traditional Chinese Medicine
TMYX: Tongmai Yangxin
cAMP: cyclic Adenosine MonoPhosphate
SAN: SinoAtrial Node

## ADDITIONAL INFORMATION

### Supplementary Figures

**Figure S1:**
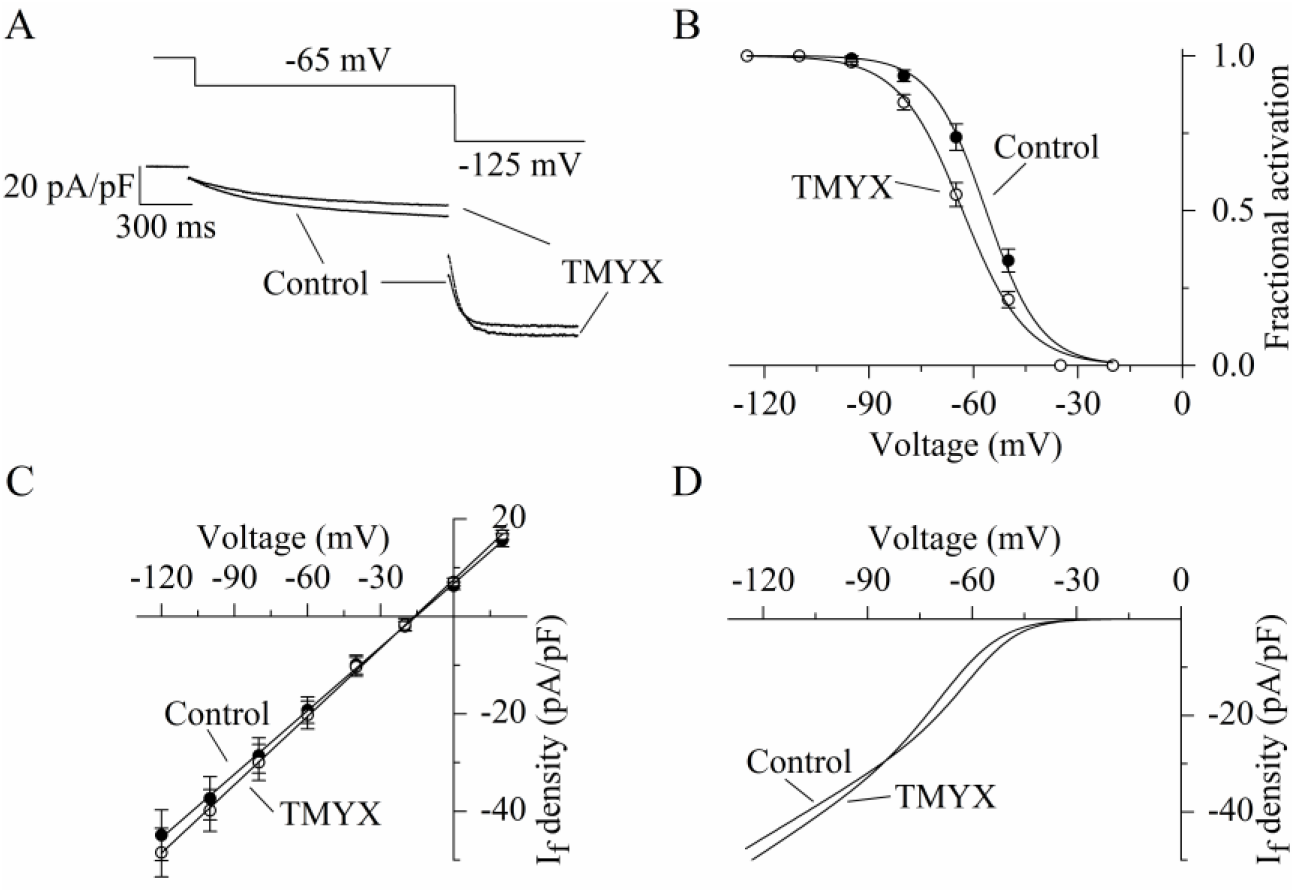
Dual action of TMYX (2 mg/ml) on the voltage-dependence and maximal conductance of the I_f_ current. A: Representative whole-cell currents (bottom) elicited by a double-step protocol (top, −65 mV/1.5 s and −125 mV/0.5 s from a holding potential of −35 mV) in control condition and during TMYX (2 mg/ml) perfusion. B: Voltage-dependent activation curves obtained in control condition and during TMYX perfusion; each data-point represents the mean±SEM fractional activation value at the voltages indicated (n=6 cells). Fitting of data by the Boltzmann equation (continuous lines) yielded half-activation values (V_½_) and inverse-slope factors (s) of −56.4 mV and 7.9 mV in control and −63.1 mV and 9.2 mV in the presence of TMYX; curves are significantly shifted (P<0.01, Extra sum-of-squares F test). C: Mean fully-activated current/voltage (I/V) relations measured before and during drug perfusion (n=6 cells). The slope of the linear regression fitting in the presence of TMYX was significantly increased (+7.5%, P<0.01, linear regression analysis test). D: Steady-state fitting currents obtained by multiplying the continuous lines shown in panels B (activation curves: Boltzmann fitting) and C (fully-activated I/V relation: linear fitting) in control condition and in the presence of the drug. At potentials more positive than the crossover point (−85 mV), the contribution of the negative shift induced by TMYX (2 mg/ml) is greater than the increment in conductance, while the opposite occurs at more negative potentials. In the pacemaker range of potentials TMYX has the net effect of reducing the contribution of the I_f_ current to the diastolic depolarization.

**Figure S2:**
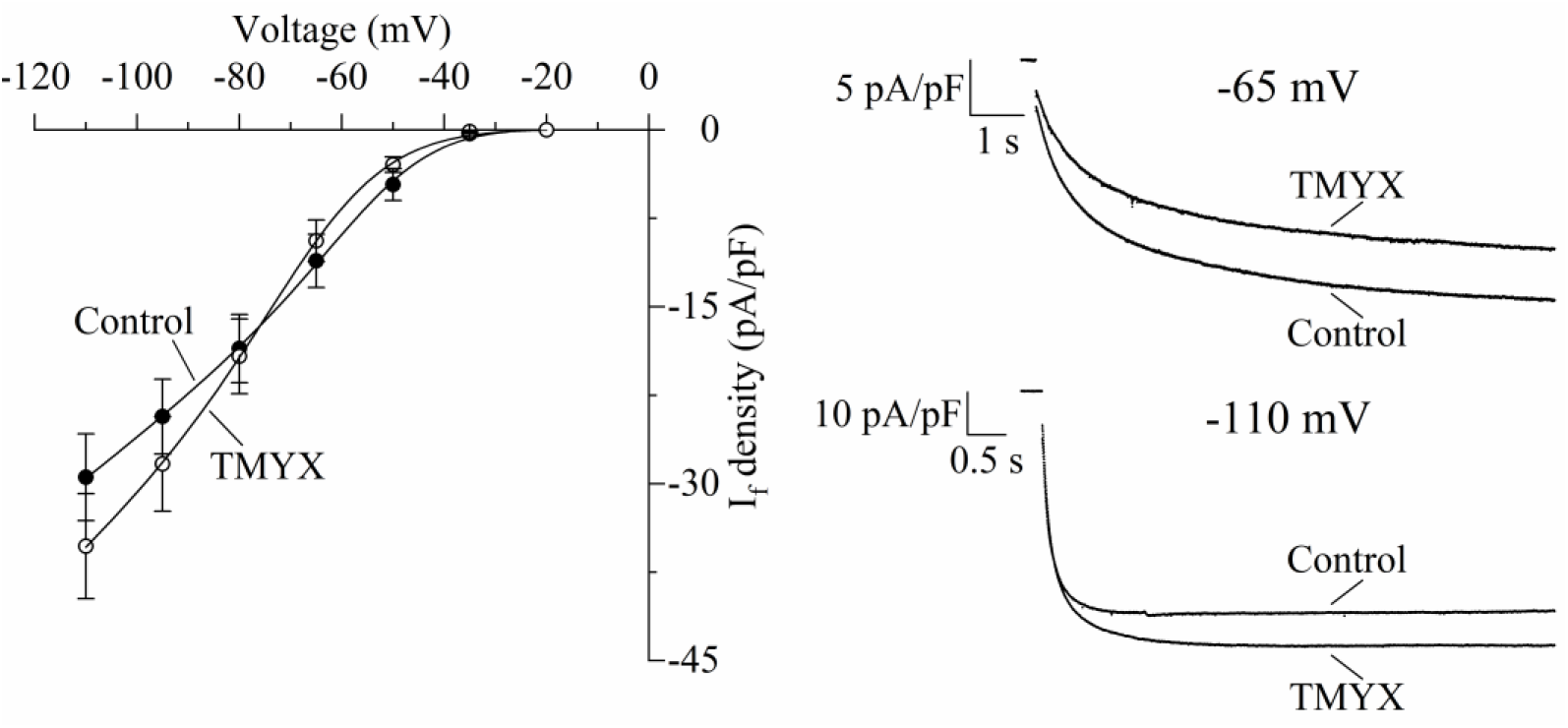
Effects of TMYX (6 mg/ml) on the voltage-dependence of the steady-state I_f_ current measured at different voltages. Left: mean±SEM steady-state current/voltage relation obtained from n=7 cells in control (filled circles) and during TMYX perfusion (empty circles). Currents were elicited by a train of 8 hyperpolarizing steps encompassing the voltage range −20/-110 mV (ΔV between steps, −15 mV); step durations were longer at more depolarized potentials to ensure attainment of complete steady-state. Continuous lines through data-points represent the best fitting procedures (obtained using the following equation: I_density_=(a*V+b)*(1/(1+exp((V-V_½_)/s))) which describes the product of a Boltzmann relation times the fully-activated I/V relation. Best fitting yielded the following values: a, 0.339 and 0.394 (pA/pF)/mV; b, 7.663 and 7.296 mV; V_½_, −51.4 and −64.3 mV; s, 10.2 and 11.5 mV, respectively for Control and TMYX conditions. Right: Representative sample traces recorded in control condition and in the presence of TMYX (6 mg/ml) at 2 different potentials (−65 mV and −110 mV). In the diastolic depolarization voltage range the net effect of TMYX is a reduction of the I_f_ current.

**Figure S3:**
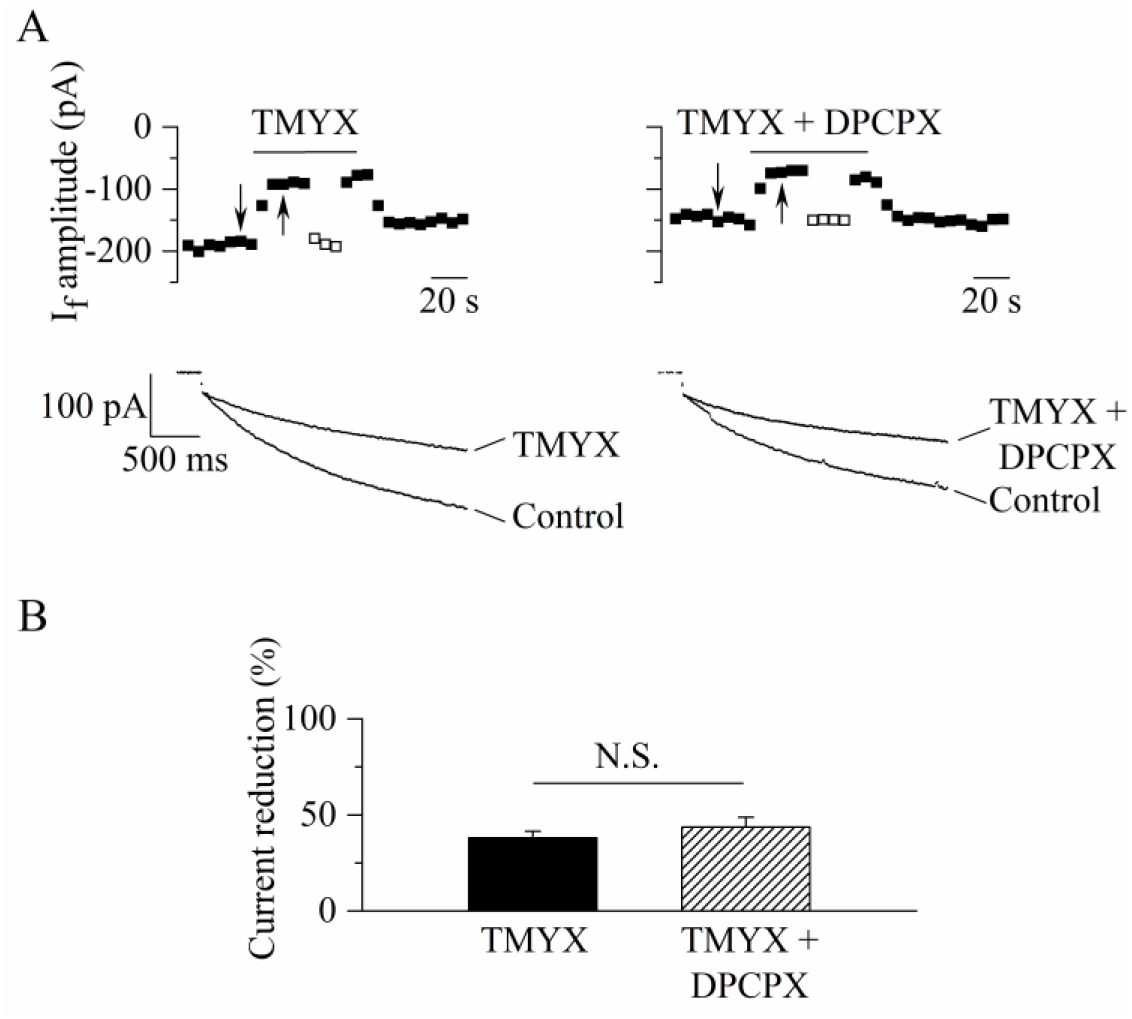
The action of TMYX is not mediated by the activation of the adenosine receptor. A: Representative steady-state time courses (top) and sample current traces (bottom) recorded at −65 mV for 2.75 s (holding potential, −35 mV) in control condition and in the presence of TMYX (6 mg/ml, left) delivered alone or in combination with the adenosine A1-receptor blocker DPCPX (Martinson, Johnson, & Wells, 1987) (1 µM, right). Empty squares represent steady-state current amplitudes recorded during manual adjustment (hyperpolarization) of the holding potential to compensate for the effect of the drugs. B: Bar-graph of the steady-state current reduction (mean±SEM %, n=6): TMYX, −38.1±3.5%; TMYX+DPCPX, −43.6±5.2%. N.S. Not Significant, P=0.111 (Student’s Pair t-Test).

**Figure S4:**
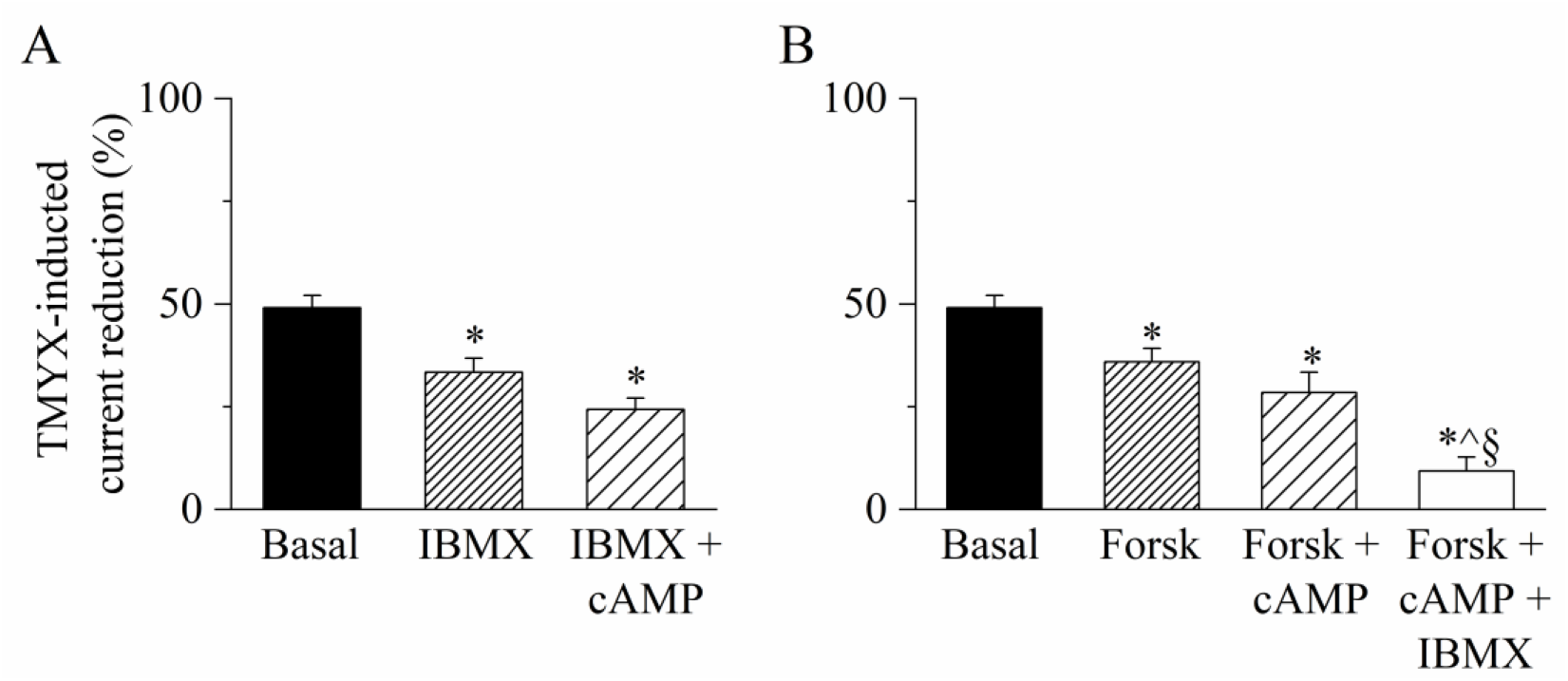
Increasing concentrations of intracellular cAMP reduce the I_f_ inhibitory action of TMYX. A: The bar-graph illustrates the effects of TMYX (6 mg/ml) on the whole-cell I_f_ current (elicited at −65 mV) assessed in Basal condition, in the presence of the phosphodiesterase inhibitor IBMX, and of IBMX+cAMP. IBMX and cAMP, (100 µM and 10 µM, respectively) were added to the pipette/intracellular solution, while TMYX was externally delivered. Mean±SEM TMYX-induced current reduction (%) values were: Basal, 49.1±3.0%, n=15; IBMX, 33.4±3.4%, n=10; IBMX+cAMP, 24.3±2.8%, n=4. *P<0.01, IBMX *vs* Basal; *P<0.01, IBMX+cAMP *vs* Basal. B: The bar-graph illustrates the effects of TMYX (6 mg/ml) on the whole-cell I_f_ current (elicited at −65 mV) measured in Basal condition, in the presence of a stimulator of adenylyl cyclase delivered alone (Forskolin) or in combination with cAMP and cAMP+IBMX. Forskolin, cAMP, and IBMX (100 µM, 10 µM, and 100 µM, respectively) were added to the pipette/intracellular solution, while TMYX was externally delivered. Mean±SEM TMYX-induced current reduction (%) values were: Basal, 49.1±3.0%, n=15; Forsk, 35.9±3.3%, n=5; Forsk+cAMP, 28.5±4.9%, n=5; Forsk+cAMP+IBMX, 9.4±3.4%, n=5. *P=0.022, Forsk *vs* Basal; *P<0.01, Forsk+cAMP *vs* Basal; *P<0.01, Forsk+cAMP+IBMX vs Basal; ^P<0.01 Forsk+cAMP+IBMX vs Forsk; §P<0.01, Forsk+cAMP+IBMX *vs* Forsk+cAMP. Statistical analysis was carried out using One-Way ANOVA followed by Fisher’s LSD post-hoc test for multiple comparisons.

**Figure S5:**
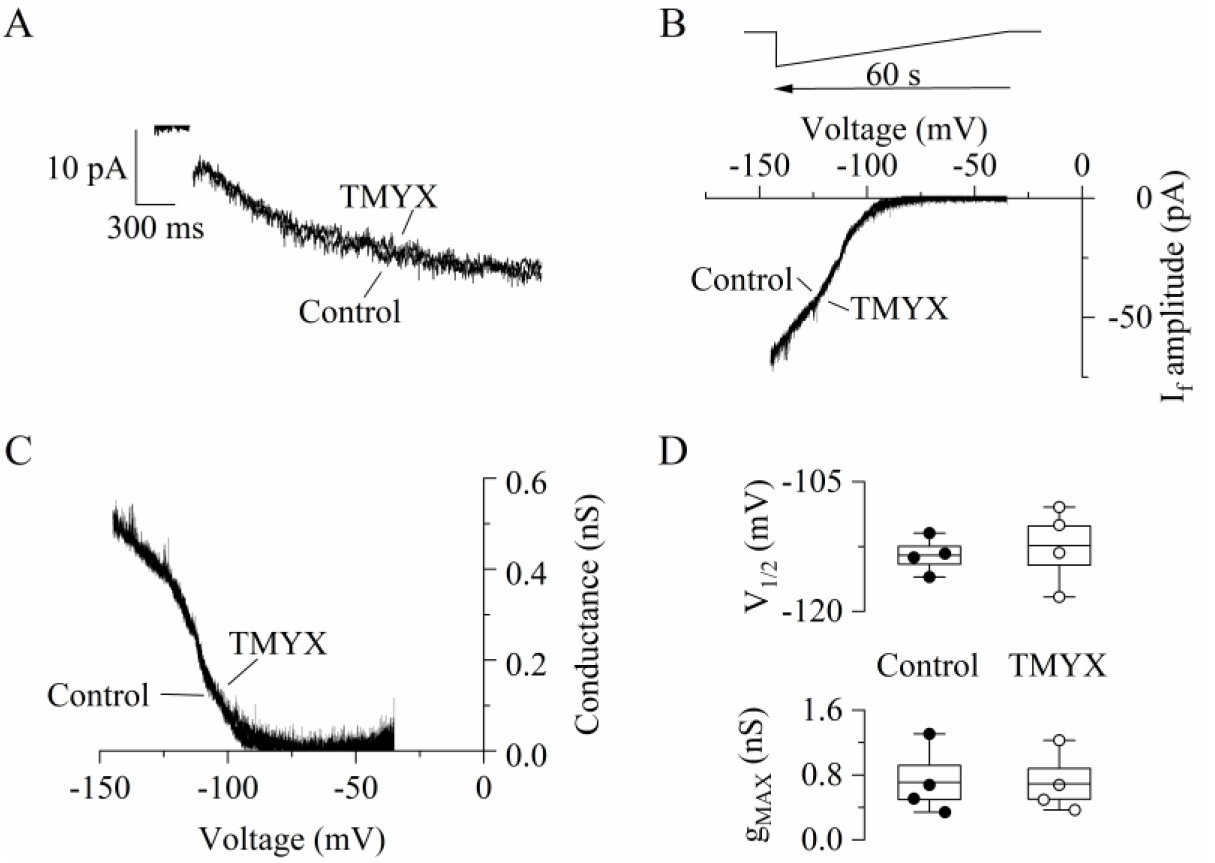
TMYX does not directly influence the intrinsic properties of f-channels. A: Representative I_f_ current traces recorded from an inside-out macropatch at a voltage of −105 mV (holding potential, −35 mV) in control condition and in the presence of TMYX (6 mg/ml). The same experiment was repeated in 3 additional patches and no significant difference in steady-state current amplitude was observed, n=4; P=0.12. B,C: Representative steady-state I/V and g/V curves obtained in an inside-out macro-patch sample by applying a slowly hyperpolarizing ramp (from −35 mV to −145 mV, rate −110 mV/min, B, top) in the absence (Control) and in the presence of the drug (TMYX). D: Box plot showing V_½_ (top) and g_max_ (bottom) values obtained from Boltzmann fitting of the g/V curves shown in panel C (n=4 patches). In the box, the middle line represents the mean value, the extremities stand for SEM and the whiskers are the maximum and minimum values. Statistical comparison of experimental data revealed that TMYX does not exert any modulatory action on these parameters (V_½_: Control, −113.5±1.0 mV; TMYX, −112.4±2.3 mV, P=0.451; g_max_: Control 0.71±0.21 NS; TMYX, 0.69±0.19 NS, P=0.528; Student’s Pair t-Test).

